# A Benchmark of Evo2 Genomic AI Models for Efficient and Practical Deployment

**DOI:** 10.1101/2025.09.10.675279

**Authors:** Huimin Li, Hongyi Ji, Yuchen Zeng, Wei Lv, Jianmin Wu, Sheng Liu, Chunhua Lin, Huanming Yang, Zhaorong Li, Yubao Chen, Wei Dong

## Abstract

The rapid advancement of DNA foundation language models has brought about a transformative shift in genomics, allowing for the deciphering of intricate patterns and regulatory mechanisms embedded within DNA sequences. The genomic foundation model Evo2 demonstrates remarkable capabilities in decoding DNA functional patterns through cross-species pretraining. However, despite the great potential of Evo2 in basic genomics research, there is currently no clear and systematic guidance on its specific application scenarios, performance, and optimization directions in the field of tumor genomics, and its performance dependency on specialized hardware (such as FP8 precision on H800 GPUs) has not been empirically benchmarked. Here, we present a focused validation of Evo2 using two independent cancer genomic datasets (Bladder Urothelial Carcinoma and Ovarian Cancer), we tested the downstream tasks of Evo2, including the prediction of tumor pathogenic variants and the prediction of mutational effects, and compared its performance on A100 and H800 GPUs. The results show that critical importance of FP8 precision, enabling the H800 to achieve a 4× faster inference speed than the A100 with stable accuracy (AUC 0.88-0.95). The 7B-parameter model emerged as the top performer, whereas the 40B model experienced a severe performance drop (AUC to 0.48) on non-FP8 hardware like the A100. These findings empirically validated Evo2’s hardware specifications and provided practical insights for researchers implementing the model with similar computational resources. Futhermore, our findings provide a framework for the application and optimization of downstream tasks of the DNA language model Evo2 in cancer, and can guide researchers in effectively applying it in genomic studies.

**Key Points:**

- Hardware Precision Impact: FP8 precision on H800 GPUs is critical for Evo 2’s performance, enabling 4× faster inference than A100 (without FP8 support) while maintaining high accuracy (AUC 0.88–0.95).
- Model Scale Optimization: The 7B-parameter model outperformed larger variants (e.g., 40B), which suffered severe accuracy drops (AUC as low as 0.48) on non-FP8 hardware, highlighting a balance between efficiency and performance.
- Practical Guidelines: We provide a framework for deploying Evo 2 in cancer genomics, including hardware recommendations, dataset curation, and downstream task optimization—valuable for researchers with varied computational resources.

## Introduction

Recent breakthroughs in deep learning have spurred the creation of large-scale genomic models capable of discerning meaningful patterns within DNA sequences. Early deep learning models like DeepBind^1^ and Basset^2^ used CNNs to predict transcription factor binding, while later models such as DNABERT^3^ and Nucleotide Transformer^4^ adopted attention mechanisms to process sequences. Specialized architectures have emerged for different tasks: Enformer^5^ excels in enhancer-promoter interaction analysis, DNA-Diffusion^6^ applies diffusion models for sequence generation, and HyenaDNA^7^ uses long-context modeling for multi-mega base sequences.

While Evo2 pre-trained on over 300 billion nucleotide sequences from a vast array of species, distinguishing itself from the traditional single-task prediction methods and emerging as a leading foundation model^8^. Evo2 utilizes masked modeling and evolutionary-scale pretraining techniques to capture universal genomic function and regulation patterns. This unique approach enables diverse applications, such as predicting variant effects, detecting regulatory elements, and conducting cross-species sequence analyses. Evo2 model stands out due to three key innovations: (1) a hierarchical attention mechanism that can capture both local motifs (like CNN-based BPNet^9^) and long-range dependencies, (2) multi-species pretraining that provides evolutionary insights beyond cross-species models like GeneCompass^10^, and (3) an efficient tokenization strategy for processing mega base-scale sequences, addressing limitations of earlier position-encoding methods in models like GPN^11^. As an evolution of the original Evo^12^ system, Evo2 advanced data management, model architecture, and training and inference capabilities, enabling accurate identification of biological units across the entire tree of life—particularly within the extensive non-coding regions of complex eukaryotes.

Evo2 has demonstrated a promising performance in predicting functional genomic elements and classifying disease-causing variants. This advancement offers an unprecedented artificial intelligence tool for applications in genetic disease diagnosis, early cancer detection, and synthetic biology. This study aims to offer a practical guide to operating Evo2, equipping researchers with the knowledge to effectively utilize this powerful tool in their genomic analyses. We walk through the step-by-step process of setting up Evo2, from installation to running initial analyses. Additionally, we will conduct a comparative analysis of Evo2’s performance on different system hardware, specifically on the H800 and A100 GPUs. By evaluating key factors such as processing speed, memory efficiency, and analytical accuracy, this study aims to identify the most suitable hardware configurations for various types of downstream analyses. The findings provide valuable guidance for researchers in selecting optimal hardware setups and offer deeper insights into Evo2’s performance, capabilities, and limitations in large-scale genomic applications.

## Materials and Methods

### Software Download and Dependency Installation

Evo2 can be obtained from its official GitHub repository (https://github.com/ArcInstitute/evo2), which provides comprehensive source code and installation guide lines. Additional dependencies for Evo2 are installed via pip or build tools as specified in the GitHub repository’s README.md. Key dependencies may incl ude deep learning frameworks (e.g., PyTorch with CUDA support), genomics libraries (e.g., Biopython), and data processing tools. Users should run the recom mended installation commands from the official repository. Alternative: If issues arise, install via the Vortex inference code following Vortex instructions. PyPI support is pending.

The computational environment configuration requires loading the following libraries and their corresponding versions to ensure compatibility with dependencies and the CUDA toolchain:

### Availability of Datasets

The official demo dataset, and runnable code provided by Evo2 are available a t the following GitHub repository: https://github.com/ArcInstitute/evo2/tree/main. The two cancer datasets utilized in this study were jointly generated by the Hangzhou Institute of Medicine and its affiliated partner hospitals. For data requests, please contact the corresponding author.

### Hardware Performance Comparison

To evaluate Evo2’s performance on different GPUs (H800 and A100), we measure:

a. Processing Speed: Time per batch for inference/training.
b. Memory Usage: GPU memory consumption during runtime.
c. Accuracy: Performance metrics (e.g., AUC - ROC) on the demo/own dataset.

Tests are conducted under identical conditions (e.g., batch size, input data, and task type), with each GPU tested sequentially in the configured environment.

## Results

We present the test datasets, experimental setups, and results of the Evo2 model on H800 and A100 platforms. We evaluated the model’s performance in pathogenicity classification and region impact analysis across three datasets: the Evo2_demo dataset (Supplementary Data 1), and two experimental datasets - Dataset1^13^ (Bladder Urothelial Carcinoma, BLCA) and Dataset 2 (ovarian cancer, OV). Further we analyzed the effects of hardware precision (FP8 support) and model scale (1B to 40B) on accuracy, computational efficiency, and result reproducibility.

### Test Datasets and Experimental Setup

Three cancer-related datasets were employed to evaluate the Evo2 model, spanning analytical objectives and cancer types (Fig 1, Supplementary Table1). The Evo2_demo (3,893 SNV, BRCA) and Dataset2 (83,401 SNV, OV) was used for pathogenicity classification, while the Dataset1 (4,791 SNV, BLCA) assessed regional impact of mutations.

**Figure 1.**
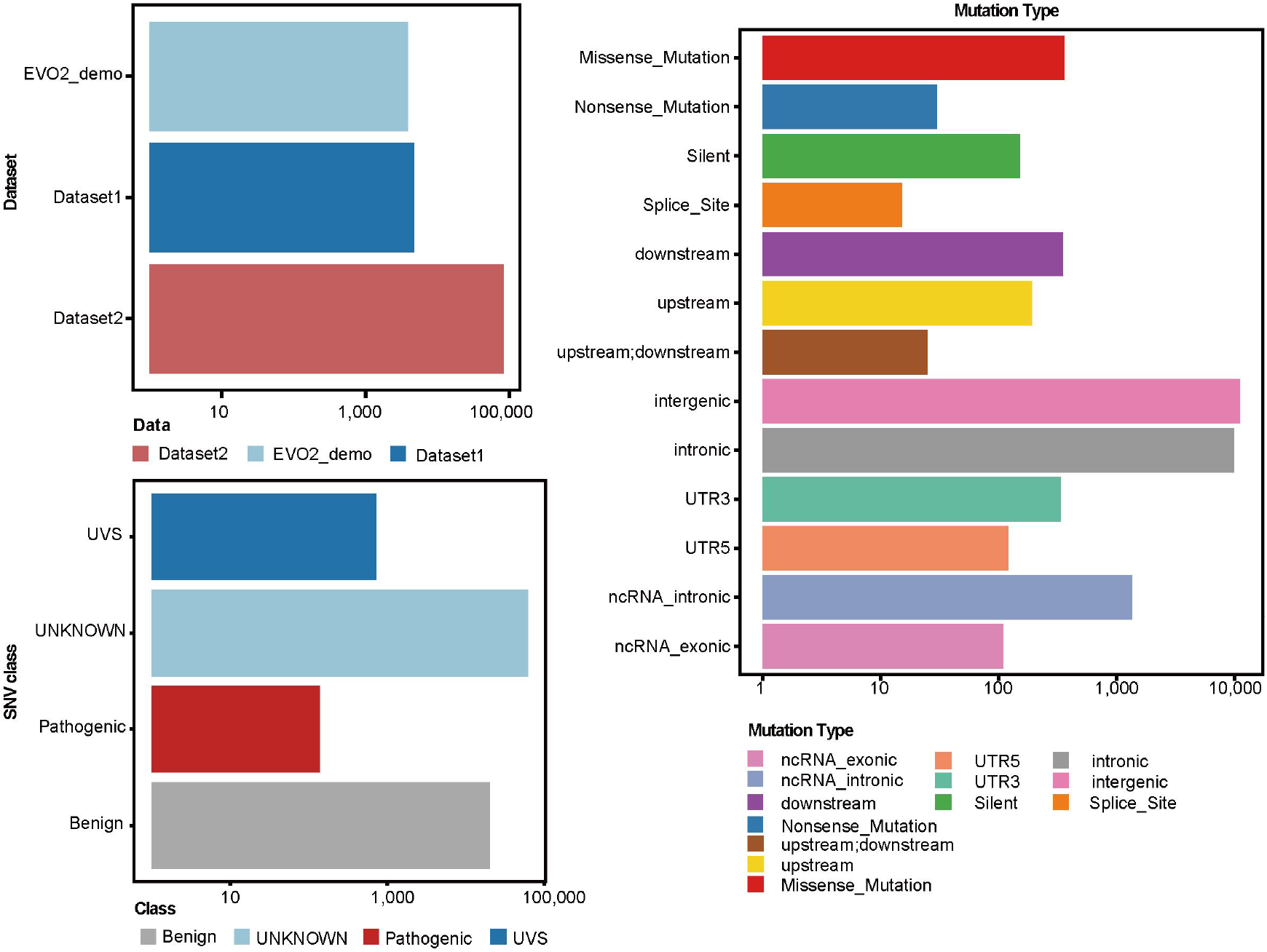
Benchmarking data structure for Evo2 variant prediction. (A) Datasets (Evo2_demo, Dataset1, Dataset2) with sample sizes (10-100,000) and (B) variant classifications (Pathogenic, Benign, UVS, UNKNOWN) in three datasets. (C) Dataset1 mutation spectrum including protein-altering (Missense/Nonsense), non-coding (intronic/intergenic), and regulatory (upstream/downstream) variants.

The Evo2_demo is derived from data officially provided by Evo2. Dataset2, the ovarian cancer data, comes from laboratory-produced data, including a total of 60 tumor samples. The raw data has undergone quality filtering and Mutect2 processing to obtain the variant sites of the samples, and the pathogenic variants of the sites have been annotated through ClinVar. Dataset1, the urothelial carcinoma data, is from Yuhuangding Hospital in Yantai, Shandong Province, consisting of 669 tumor samples, which is the largest Asian cohort of urothelial carcinoma. The raw data have been subjected to quality filtering and Mutect2 processing to obtain the variant sites and regional information of the samples. Experiments were conducted on two hardware platforms: H800 (with FP8 precision support) and A100 (lacking FP8), to compare model performance, computational efficiency, and result reproducibility.

### Pathogenicity Classification Performance

#### H800 Platform Results

On the H800, the Evo2 model demonstrated strong consistency with official benchmarks. Evo2 zero-shot evaluation provides an initial measure of a model’s inherent ability to predict functional effects, however, different models exhibit varying predictive performances, and the 7B model achieved the best results (Fig 2B, Supplementary Fig1). For Evo2_demo dataset, the predictive performance of the Evo2 7B model for pathogenic variant sites reached to an AUC of 0.87 (Fig 2C). Larger models (40B_base/40B) showed comparable performance (AUC 0.86 - 0.87), indicating minimal benefit from increasing model size beyond 7B parameters. The 7B model correctly classified mutations (Supplementary Fig2B), with clear separation between benign and pathogenic variants compared with another model.

**Figure 2.**
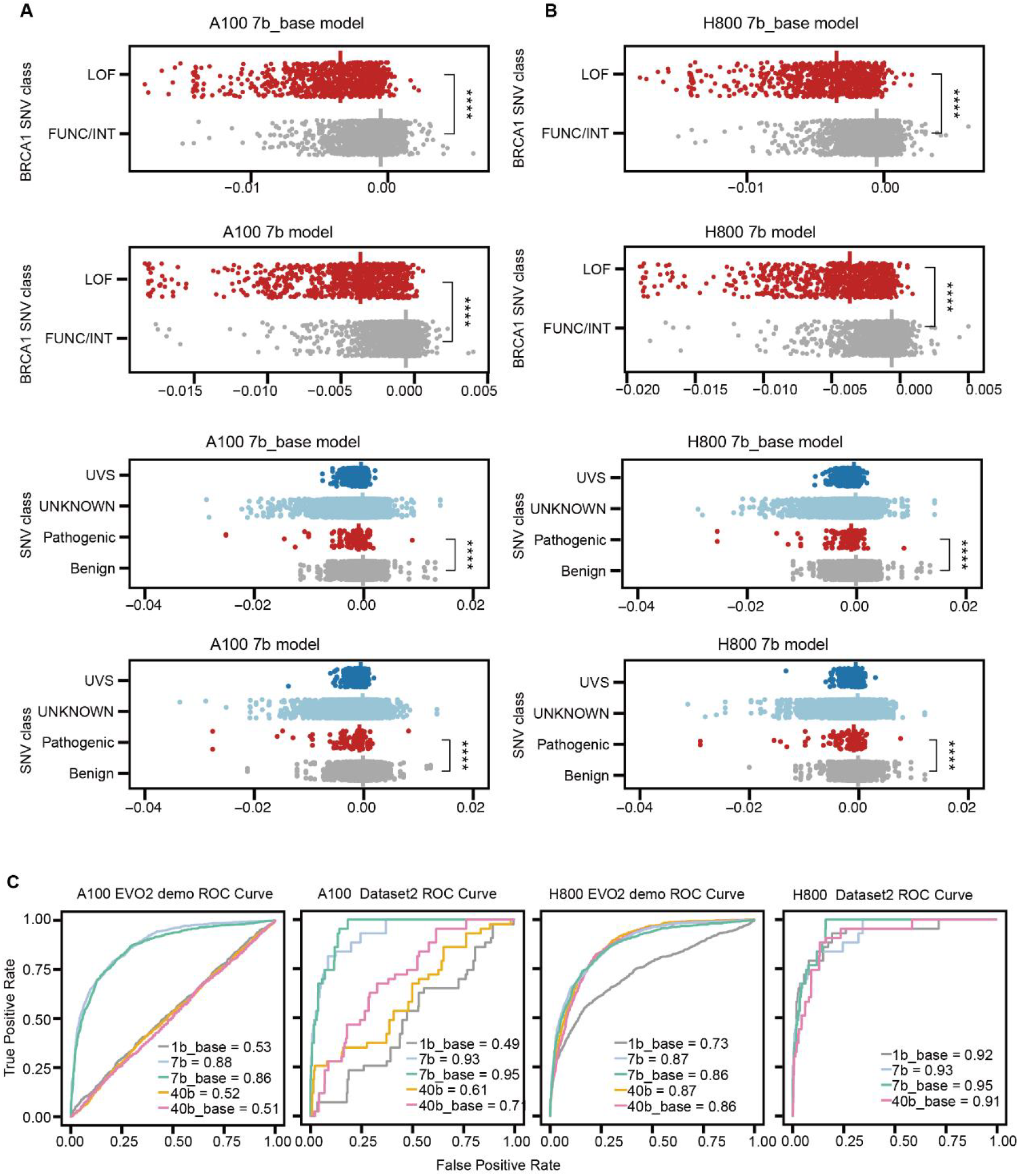
Benchmarking Evo2’s zero-shot variant scoring on A100 vs H800 GPUs. **(A)** Evo2 zero-shot likelihood scores plotted for demo and Dataset2 variants using A100, demonstrating the model’s ability to separate these classes. p-value calculated by two-sided Wilcoxon rank sum test. **(B)** Evo2 zero-shot likelihood scores plotted for demo and Dataset2 variants using H800. p-value calculated by two-sided Wilcoxon rank sum test. **(C)** AUC (Area Under the Curve) measures the model’s ability to distinguish pathogenic variants. Higher AUC (closer to 1) indicates better discrimination performance.

In the Dataset2, the 7B_base model outperformed all others with an AUC of 0.95, while the 7B model achieved an AUC of 0.93 (Fig 2C). Compared with the prediction performance of BRCA mutations on the demo data, no obvious distinction between pathogenic and non-pathogenic variants. We hypothesized that it is necessary to train different prediction models for different genes (Supplementary Fig3B).

#### A100 Platform Results

A100 performance was compromised by the lack of FP8 precision. For Evo2_demo, 7B_base and 7B models retained acceptable AUC values (0.86 and 0.88), but 40B-series models suffered a drastic drop (AUC 0.51 - 0.52), likely due to numerical instability (Fig 2C). In Dataset2, the 7B_base model achieved an AUC of 0.95 (Fig 2C), with a prolonged runtime of 28 hours and 40 minutes (Fig 3B).

**Figure 3.**
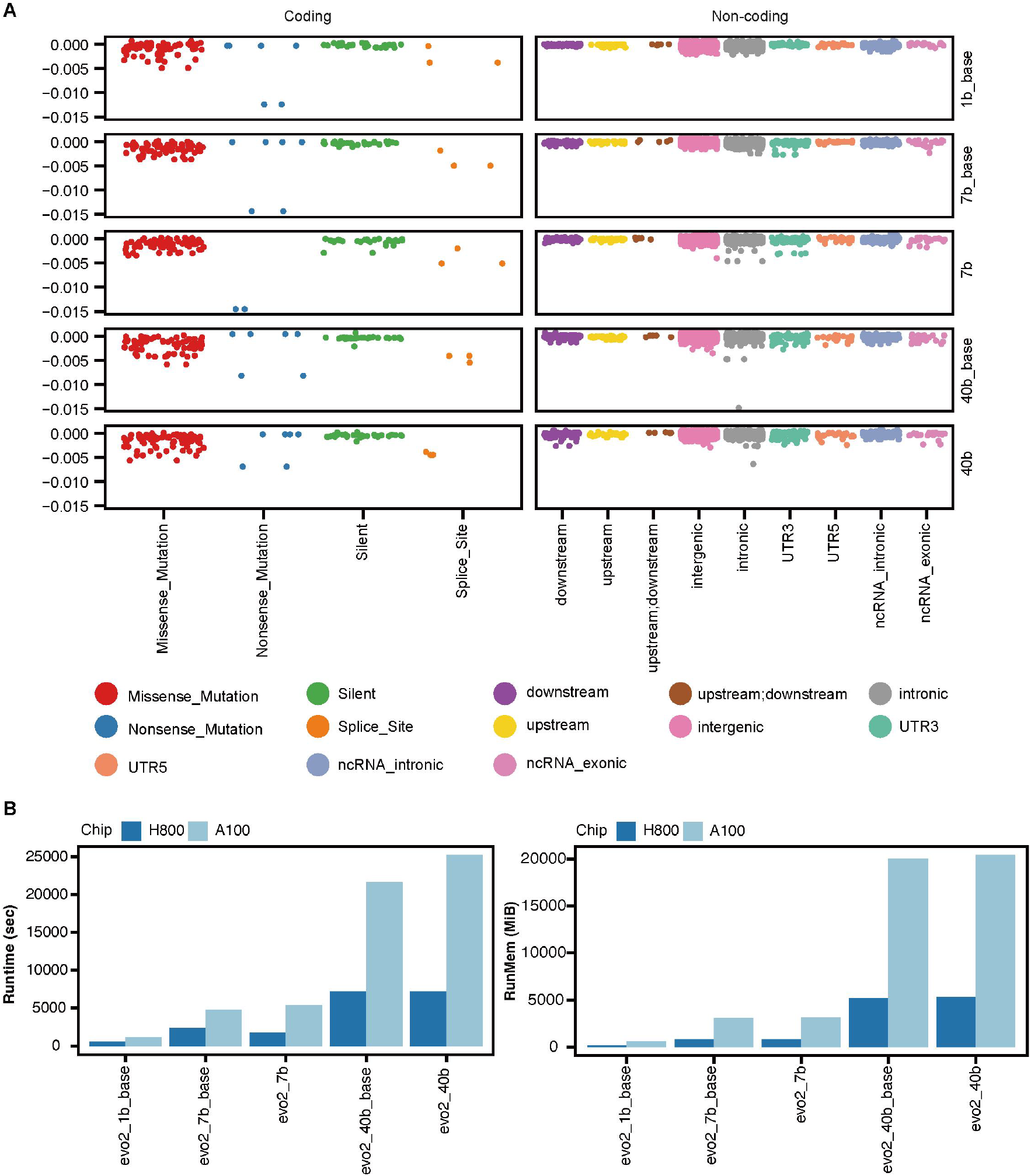
Variant type distribution and computational efficiency benchmarking. **(A)** Distribution of Genetic Variants Across Coding and Non-coding Genomic Regions by Mutation Type. Distribution of genetic variants in coding (left) and non - coding (right) regions. Colors represent variant types. Columns show different genomic context resolutions. Y - axes reflect variant - associated metric values. **(B)** Bar plots represent the runtime (in seconds) (left) and memory usage (in Mebibytes) (right) for the Evo2_demo across different models running on the H800 GPU and A100 GPU.

### Pathogenicity Classification supervised classification model

We also attempted to optimize the 1B model for pathogenic variant prediction and achieved promising results. For BRCA SNVs, we again parsed sequences of an 8,192 bp window around the mutation site using the human reference genome. We extracted embeddings from the layer of final block of the Evo2 1B model for the reference sequence and the SNV, which were used as inputs to train a classification model that consists of a feed-forward neural network with one hidden layer. The neural network took an input of dimension and processes it through fully connected layers of sizes 1920 neurons, and outputting a probability that given SNV is pathogenic.

### Region Impact Analysis

On H800, all Evo2 variants distinguished mutation score distributions between coding and non-coding regions (Fig 3A), demonstrating the model’s ability to capture functional impacts. In contrast, A100 models failed to differentiate these regions (Fig 3A), highlighting the critical role of FP8 precision in fine-grained feature analysis.

### Computational Efficiency and Resource Usage

H800 outperformed A100 in speed and memory efficiency. For example, the 7B model processed Evo2_demo in 1.5 hours with 3,156 MiB on H800, versus 6 hours and 20,024 MiB on A100 (Fig 3B). Larger models (e.g., 40B) showed disproportionate memory usage (20,000+ MiB) without performance gains, underscoring the trade-off between model scale and usability.

Single-sample inference on H800 was highly efficient: 0.13 seconds/sequence for 1B_base and 1 second/sequence for 40B (Fig 3B), enabling real-time applications for smaller models.

## Discussion

Our evaluation demonstrates that Evo2 optimized for FP8 precision on the H800 platform delivers robust performance on genomic datasets. Notably, the 7B_base model achieves an optimal balance between predictive accuracy (AUC = 0.95) and computational efficiency (0.42 s/sequence), positioning it as an ideal solution for large-scale biomedical applications. Hardware benchmarking confirms FP8 acceleration as a cornerstone for the platform’s computational efficiency. The significant performance gap between H800 and A100 architectures, particularly with larger models, underscores the critical need to prioritize H800-based FP8 optimization for scalable handling of complex multimodal biological data.

However, although the model showed strong predictive performance for pathogenic variants on the demo dataset, its performance on the independently collected test data is suboptimal. We speculate that the reason may be either the insufficient number of annotated variant types in the data or the excessive variety of variant genes, making the model unable to effectively evaluate pathogenic variants other than BRCA genes. If it is to be used for the assessment of pathogenic variants, further optimization of the model is required. The Evo2 paper mentions that the embedding values obtained from the model can be used to construct unsupervised models for specific downstream tasks. We also attempted to optimize the 1B model for pathogenic variant prediction and achieved promising results. Therefore, we recommend collecting more data to build a tool capable of detecting multi-gene pathogenic variants.

Building on these remarkable strengths, we first envision Evo2 as the cornerstone of a unified digital research platform tailored for multimodal biological studies. This platform’s core value lies in its ability to leverage Evo2’s high - performance genomic analysis capabilities while integrating shared multi - omics tumor data, high - performance computing resources, and scalable storage infrastructure. Such integration is set to accelerate discoveries in precision medicine across a wide range of domains. These domains include multi - omics oncology, where it can dissect complex tumor ecosystems; rare disease genomics, aiding in the identification of causal variants; clinical risk stratification, enabling more accurate patient prognosis; and GWAS analysis, facilitating the exploration of genetic associations with diseases.

We invite the research community to leverage this powerful platform to drive transformative discoveries in precision medicine. We welcome collaborations with academic institutions, clinical researchers, and pharmaceutical partners to further develop and apply this technology to pressing challenges in genomics and precision health. Together, we can harness the full potential of AI-driven genomic analysis to advance human health.

## Supplementary Data

Supplementary tables and figures are attached at the end.

## Author contributions

Huimin Li: Data curation, formal analysis, validation, investigation, visualization, methodology, writing original draft, writing-review, and editing. Hongyi Ji: Formal analysis, writing original draft, investigation, visualization, writing review and editing. Yuchen Zeng: Data curation, formal analysis, validation, investigation, visualization, writing original draft, writing review and editing. Wei Lv: Data curation, writing-review and editing. Jianmin Wu: Data curation, writing-review, and editing. Sheng Liu: writing-review and editing, investigation. Chunhua Lin: Data curation. Huanming Yang: Resources, supervision, investigation, writing-review, and editing. Zhaorong Li: Conceptualization, resource, supervision, investigation, writing-review, and editing. Yubao Chen: Resources. Wei Dong: Conceptualization, Resources, supervision, investigation, writing-review, and editing

## Funding

We sincerely thank the New Quality Productive Forces Center, Zhuhai Technology Industry Group for providing essential computational resources and technical support, which were pivotal to the success of this work. Their high-performance computing infrastructure significantly accelerated our research progress.

---

**Table 1.** Software and Dependency Versions for Genomic Deep Learning.

| Packages/Library | Version |
| --- | --- |
| Python | 3.12 |
| CUDA | 12.1 |
| cuDNN | 9.8 (based on CUDA 12) |
| GCC | 10.5 |
| OpenBLAS | 0.3.20 (based on GCC 10.5) |
| CMake | 3.27.0 |
| CCache | 4.10.2 |

**Table S1.** Summary of Genomic Datasets for Cancer Mutation Analysis.

| Dataset | Source | Mutation<br>Number | Cancer Type | Analysis Type |
| --- | --- | --- | --- | --- |
| Evo2_demo | Evo 2 | 3,893 | BRCA | Pathogenicity<br>Classification |
| Dataset1 | CUGA | 4,791 | BLCA | Region Impact Analysis |
| Dataset2 | HIM | 83,401 | OV | Pathogenicity<br>Classification |

**Table S2.** Model Training/Inference Performance on H800 andA100 GPUs.

| Model | H800 Runtime<br>(sec) | H800 Memory<br>(MiB) | A100 Runtime<br>(sec) | A100 Memory<br>(MiB) |
| --- | --- | --- | --- | --- |
| 1B_base | 1,200 | 612 | — | — |
| 7B_base | 4,800 | 3,108 | — | — |
| 7B | 5,400 | 3,156 | 21,600 | 20,024 |
| 40B_base | 21,600 | 20,024 | 25,200 | 20,414 |
| 40B | 25,200 | 20,414 | — | — |

**Figure S1.**
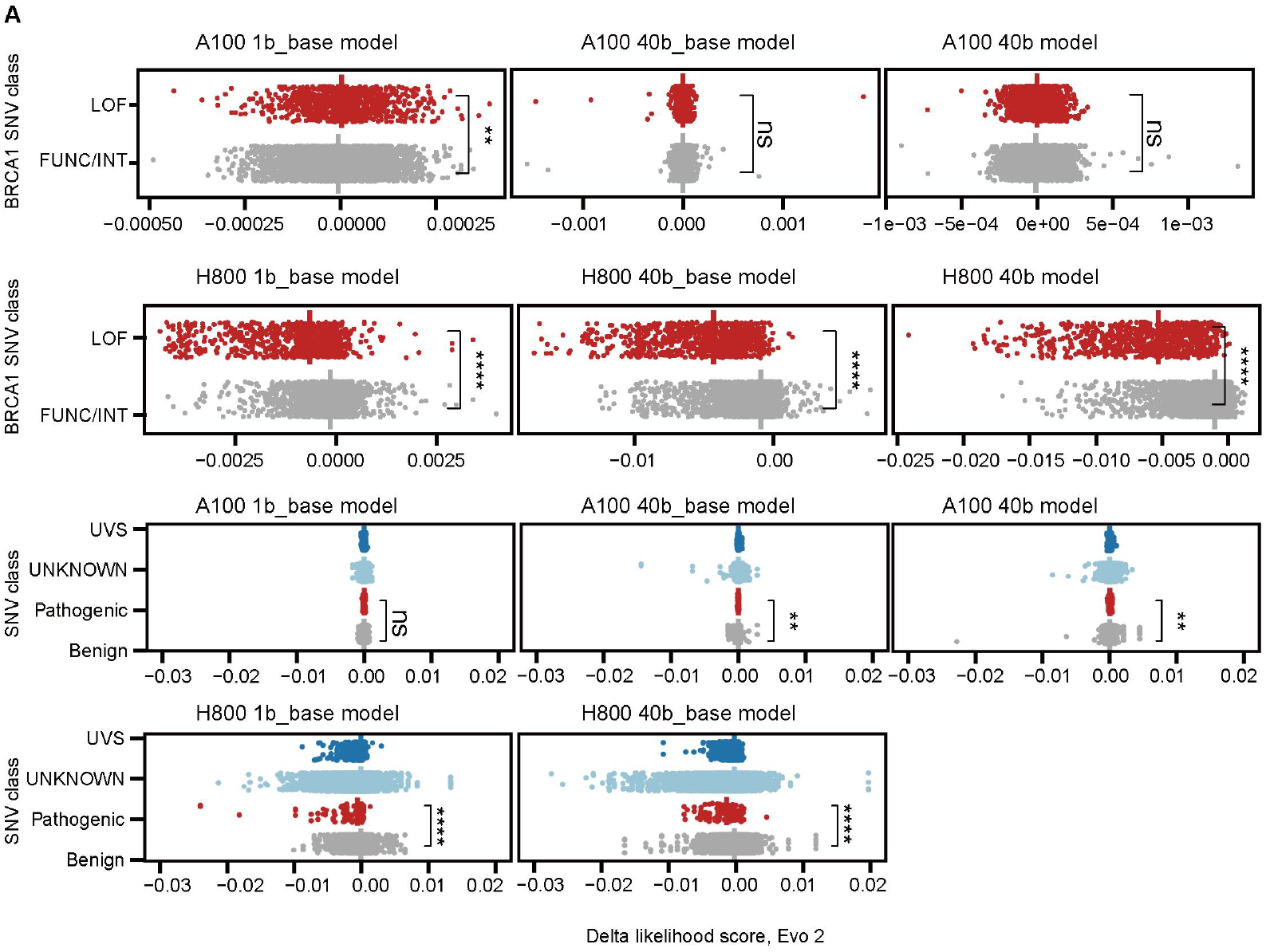
Architecture and hardware effects on Evo 2 variant scoring performance. Evo 2 zero-shot likelihood scores plotted for demo and dataset2 variants using A100 and H800. p-value calculated by two-sided Wilcoxon rank sum test.

**Figure S2.**
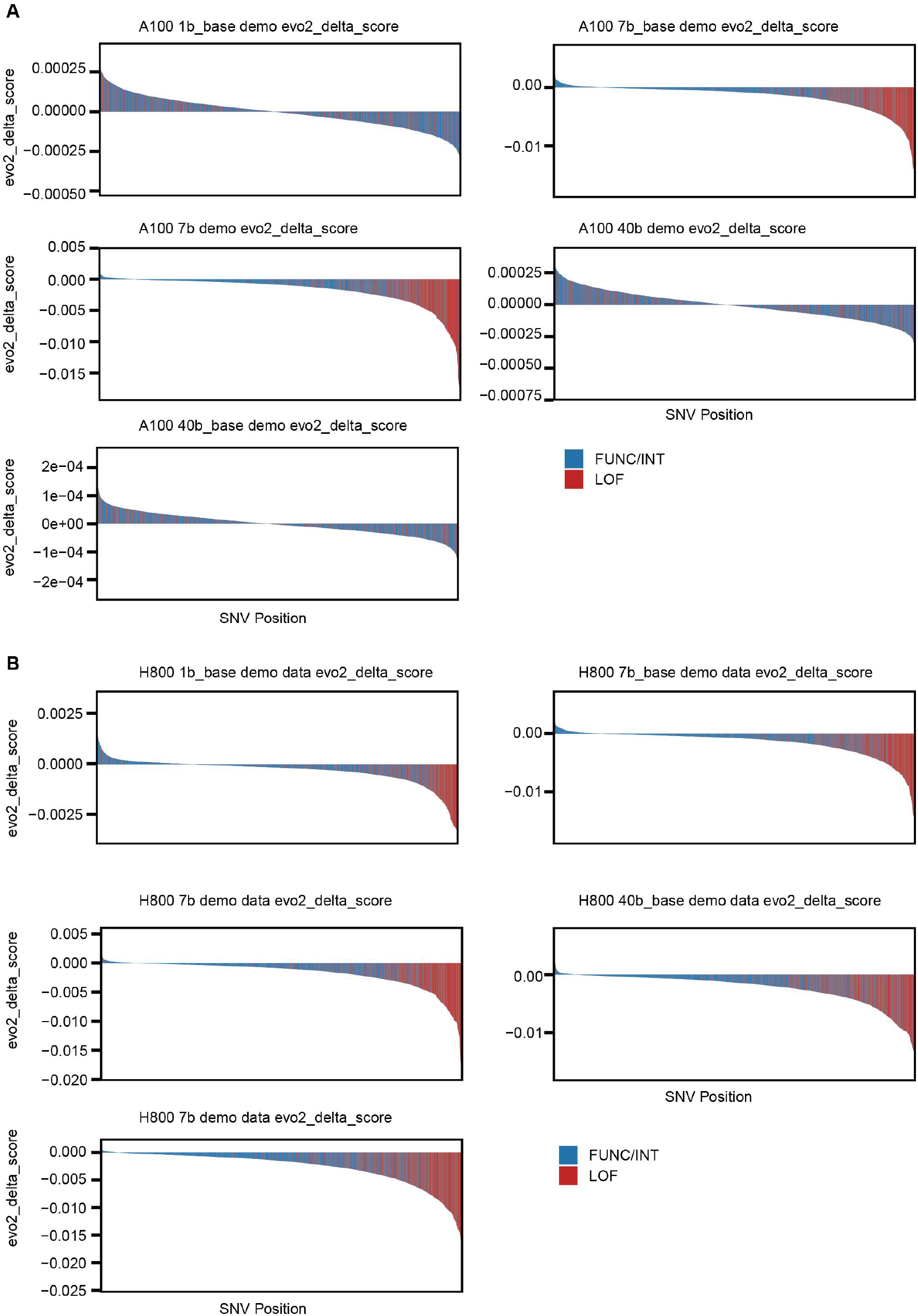
Evo 2 zero-shot likelihood delta scores in different plat. Bar plot shows the different score between FUNC/INT Mutations and LOF Mutations in demo dataset.

**Figure S3.**
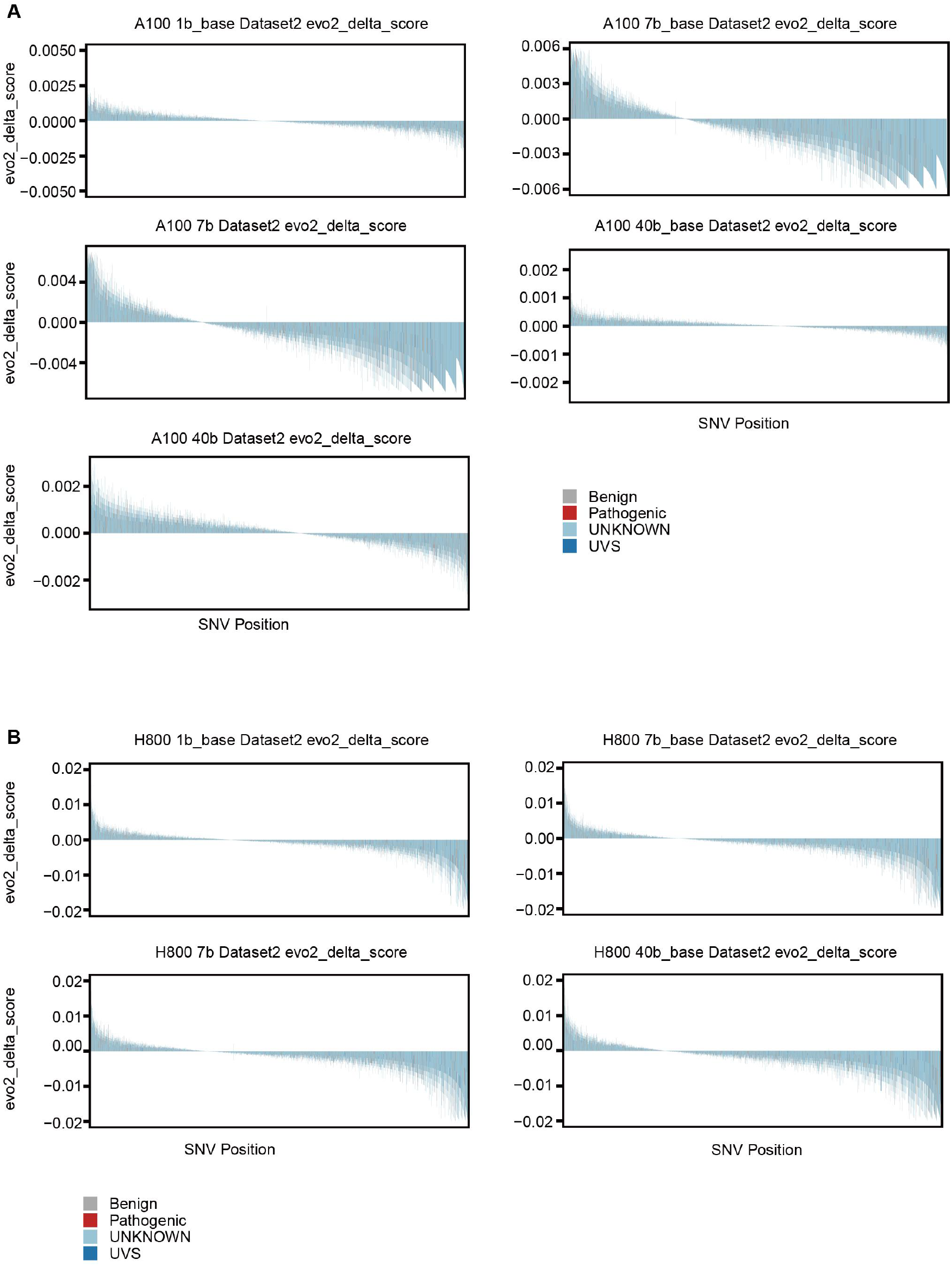
Evo 2 zero-shot likelihood delta scores in different plat. Bar plot shows the different score between FUNC/INT Mutations and LOF Mutations in Dataset2.

## Notes

### Competing Interest Statement

The authors have declared no competing interest.

